# Reactive oxygen species suppress phagocyte surveillance by oxidizing cytoskeletal regulators

**DOI:** 10.1101/2024.01.31.578237

**Authors:** Iuliia Ferling, Steffen Pfalzgraf, Lea Moutounet, Lanhui Qiu, Iris Li, Yuhuan Zhou, Sergio Grinstein, Spencer A. Freeman

**Affiliations:** Program in Cell Biology, The Hospital for Sick Children, Toronto, ON M5G 0A4, Canada; Department of Biochemistry, University of Toronto, Toronto, ON M5S 1A8, Canada

**Keywords:** NADPH oxidase, Fc-gamma receptors, phosphatidylserine, efferocytosis, Rho GTPases, ERM proteins

## Abstract

Despite their superficial similarities, the phagocytosis of pathogens differs from that of apoptotic cells in their recognition mechanisms and downstream signaling pathways. While the initial stages of these processes have been studied, the cytoskeletal reorganization that follows particle uptake is not well understood. By comparing the uptake of phosphatidylserine (PS)- coated targets versus IgG-opsonized targets of identical size, shape, and rigidity, we noted remarkable differences in the accompanying changes in cell morphology, adhesion and migration that persisted long after phagocytosis. While myeloid cells continued to survey their microenvironment after engulfing PS-coated targets, the uptake of IgG-opsonized targets caused phagocytes to round up, decreased their membrane ruffling, and led to the complete disassembly of podosomes. These changes were associated with increased activation of Rho and a concomitant decrease of Rac activity that collectively resulted in the thickening and compaction of the cortical F-actin cytoskeleton. Rho/formin-induced actin polymers were fastened to the membrane by their preferential interaction with Ezrin-Radixin-Moesin (ERM) proteins, which were necessary for cell compaction and podosome disassembly following ingestion of IgG-coated particles. The source of the distinct responses to PS- versus IgG-targets was the differential activation of the respiratory burst mediated by the NADPH oxidase: reactive oxygen species (ROS), emanating from phagosomes containing IgG-opsonized targets – but not those containing PS-coated ones – directly led to the activation of Rho. Similar findings were made with phagocytes that encountered pathogens or microbial-associated molecular patterns (MAMPS) that instigate the activation of the NADPH oxidase. These results implicate a connection between sensing of harmful particulates, the oxidation of cytoskeletal regulators, and the immune surveillance by myeloid cells that have potentially important consequences for the containment of pathogens.

## Introduction

Innate immunity serves as the first line of defense against invading pathogens and altered self-cells, playing a crucial role in maintaining homeostasis throughout the organism^1, 2^. Central to this defense mechanism are myeloid cells like macrophages, dendritic cells, and granulocytes. These cells deploy complex cytoskeletal rearrangements to recognize and eliminate potential threats^3^. Even in the steady state, myeloid cells constantly survey their surroundings for potential targets by forming ruffles and extending pseudopodia. These extensions aid in random patrolling locomotion as well as in directed chemotaxis. Some of the ruffles fuse and seal to form macropinosomes. Others serve as the first points of contact between the immune cell and phagocytic targets, which can then be sequestered into vacuolar compartments by receptor-mediated phagocytosis. In order to stabilize directional migration or initiate transcellular diapedesis, myeloid cells assemble and engage podosomes, complex actin structures that enable adhesion to the substrate and degradation of the extracellular matrix^4–6^

Having encountered their prey, myeloid cells must promptly adapt their behavior in response to the specific nature of the target engaged. While superficially similar, the recognition mechanisms and intracellular signaling pathways underlying the phagocytosis of pathogens and of apoptotic cells (efferocytosis) differ markedly. Ingested pathogens often trigger inflammation by activating endomembrane receptors or cytosolic sensors that initiate inflammasome formation^7, 8^; as a result, cytokines are released that direct the migration of additional immune cells to best combat the infection. In contrast, apoptotic targets have anti-inflammatory effects^9^.

Fcγ receptor-mediated phagocytosis is an effective means of eradicating invading pathogens: IgG-coated pathogens are thereby engulfed and eliminated within phagolysosomes. An appropriate residence time of myeloid cells in the infected area is critical. Premature migration of phagocytes carrying incompletely digested pathogenic cargo can release cytokines that attract supporting immune cells to uninfected areas and may trigger excessive^10^, unproductive inflammation. Moreover, some intracellular pathogens survive within the phagocyte. In these instances the persisting microorganisms could be transported to different tissues and organs. This dangerous scenario can give rise to secondary infection sites, exacerbating the overall illness^11, 12^. Thus, the resumption of phagocyte mobility and surveillance activity after target ingestion must be carefully timed.

Remarkably, the effect of target engagement on the exploratory and chemotactic activity of phagocytes has not been directly investigated. We used a combination of imaging and biophysical approaches to address these questions. Because the timing of reinitiation of surveillance would affect differently inflammatory and anti-inflammatory targets, we compared the effects of IgG-opsonized targets to those of phosphatidylserine (PS)-coated targets, used to mimic the engagement of apoptotic cells. Our findings implicate a profound connection between the recognition of harmful particulates, the oxidative modification of cytoskeletal regulators, and the timing of resumption of immune surveillance.

## Results

### Cytoskeletal remodeling of myeloid cells after their engulfment of PS- versus IgG-coated targets

While the remodeling of the cytoskeleton that facilitates phagocytosis has been well studied, the longer-term consequences of phagocytosis on cell dynamics have been largely neglected. We initiated our studies by evaluating the long-term responses of macrophages to target particles of identical size, shape, and stiffness coated with either 5% PS (complemented with other phospholipids) to trigger efferocytosis^13^, or with IgG to instigate Fc-mediated phagocytosis. We challenged bone marrow-derived macrophages (BMDM) with the particles and allowed an initial 10 min period of uptake, ensuring a similar number of targets were phagocytosed in both cases. Unbound particles were removed and the macrophages incubated for an additional 30 min before analysis. Scanning electron microscopy revealed clear differences in the morphology of the BMDM after uptake of PS- versus IgG-coated targets (**Fig 1a**). BMDM appeared to contract and round up after ingestion of IgG-coated particles. In stark contrast, BMDM that had ingested PS-coated beads retained their spread morphology and broad dorsal ruffles (**Fig 1a**). Contraction was quantified by measuring the area of the cells in contact with the coverglass, monitoring the distribution of F-actin by confocal microscopy using fluorescent phalloidin. These estimates confirmed the differential behavior of the cells following ingestion of the two types of beads (**Fig 1b**).

**Figure 1.**
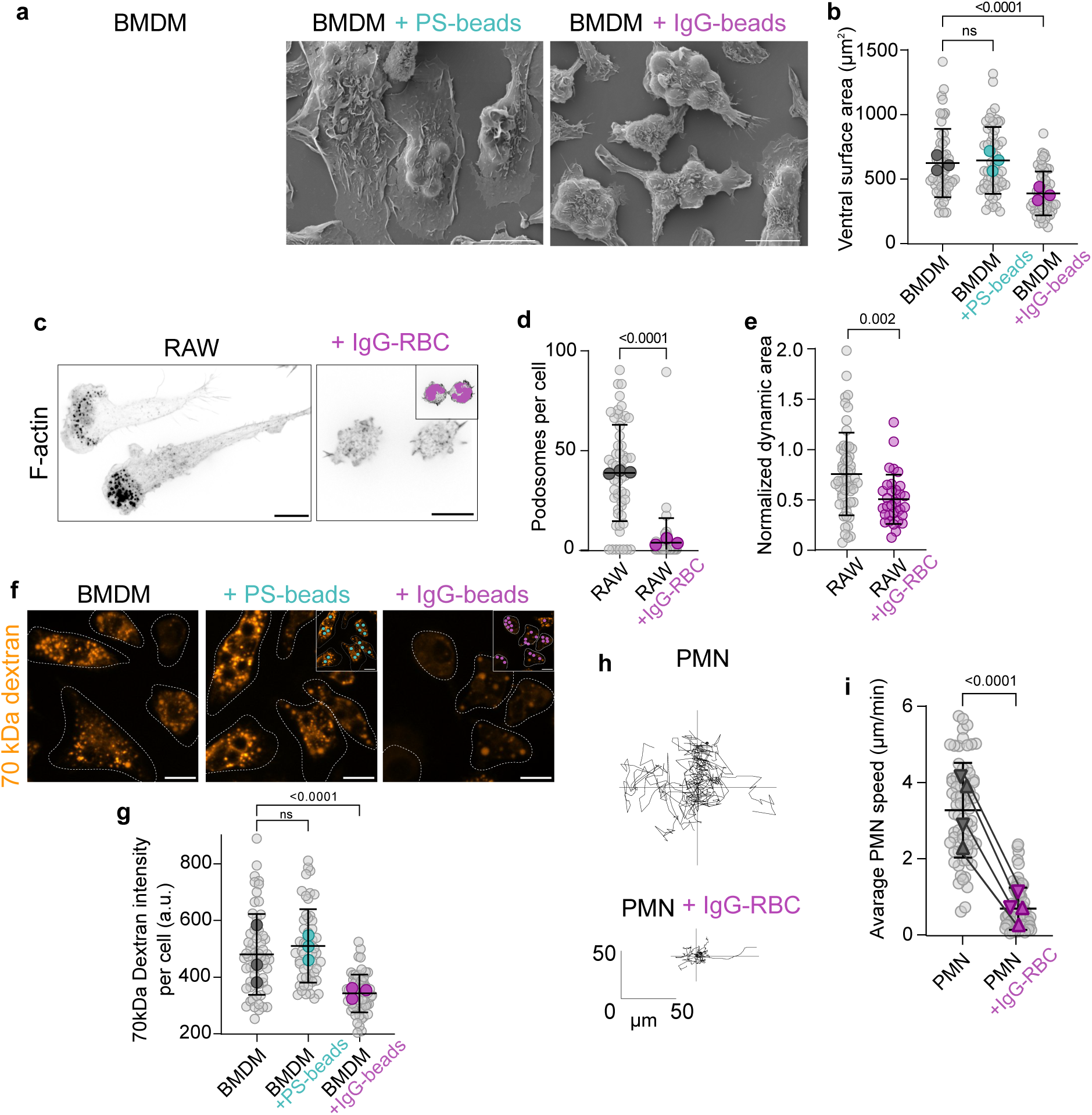
Phagocytosis alters immune surveillance. **a)** Scanning electron micrographs of bone-marrow-derived macrophages (BMDM) 30-40 min after phagocytosis of 4 μm polystyrene microspheres coated with phospholipids including 5% phosphatidylserine (PS) or opsonized with IgG. **b)** Quantification of the ventral surface area of BMDM, determined by visualizing F-actin at the level of cell attachment to the coverslip. Only BMDM containing ≥3 particles were quantified. Here and elsewhere grey dots represent individual cells and colored dots represent means of separate experiments. n = 3. **c-d)** RAW264.7 cells were stained for F-actin before or 30-40 min after phagocytosis of IgG-opsonized sheep red blood cells (RBC). Representative confocal ventral images are shown in **c**. Inset, extended focus showing IgG-RBC (magenta). In **d**, the number of podosomes were quantified; cells from 3 independent experiments. **e)** Normalized ruffling cell area estimated over 10 min by acquiring images of LifeAct-expressing RAW 264.7 cells every 15 s as described in Methods and in S Fig 2. Data from 3 independent experiments. See also *S Video 1*. **f-g)** Uptake of 70 kDa TMR-dextran (orange) by BMDM over 10 min before and after the uptake of the indicated particles, shown in insets. Panel **g** shows quantification of dextran uptake in cells from 3 independent experiments. **h-i)** Chemokinesis of human neutrophils stimulated with 10 μM fMLP before and after phagocytosis of IgG-RBC. Trajectories from one video, tracking single cells every 10 s for 10 min. See also *S Video 2*. In **i** grey dots represent the average speed of human neutrophils from heathy donors; ▴ = female donor; ▾ = male donor. *P* values are determined using a one-way ANOVA **(b, g)** or Student’s *t*-test **(d, e, i)**. All scale bars, 10 μm

More detailed analysis of the distribution of phalloidin-stained cells also revealed that prominent F-actin-dense foci, which we subsequently identified as podosomes (*see below*), were entirely disassembled in cells that had ingested IgG-coated particles (**Fig 1c-d**). Dynamic membrane ruffling, determined measuring the surveillance area of RAW 264.7 cells expressing actin-GFP by video recording (*see* **Video 1, S Fig 2** *for details*), was also depressed in cells that had ingested IgG-opsonized targets (**Fig 1e**). Membrane ruffles formed by macrophages can fold back on the cell body and fuse, resulting in the formation of macropinosomes that entrap the surrounding fluid medium^2^. We measured the uptake of labeled 70 kDa dextran, which is internalized preferentially by macropinocytosis, as an alternate means of assessing ruffling activity. Convincingly, we found that dextran uptake decreased considerably in BMDM that had ingested IgG-opsonized targets, but not PS-targets (**Fig 1f-g**). Together, these findings imply that Fc-mediated phagocytosis –but not efferocytosis– had a marked impact on actin remodelling and membrane dynamics.

**Figure 2.**
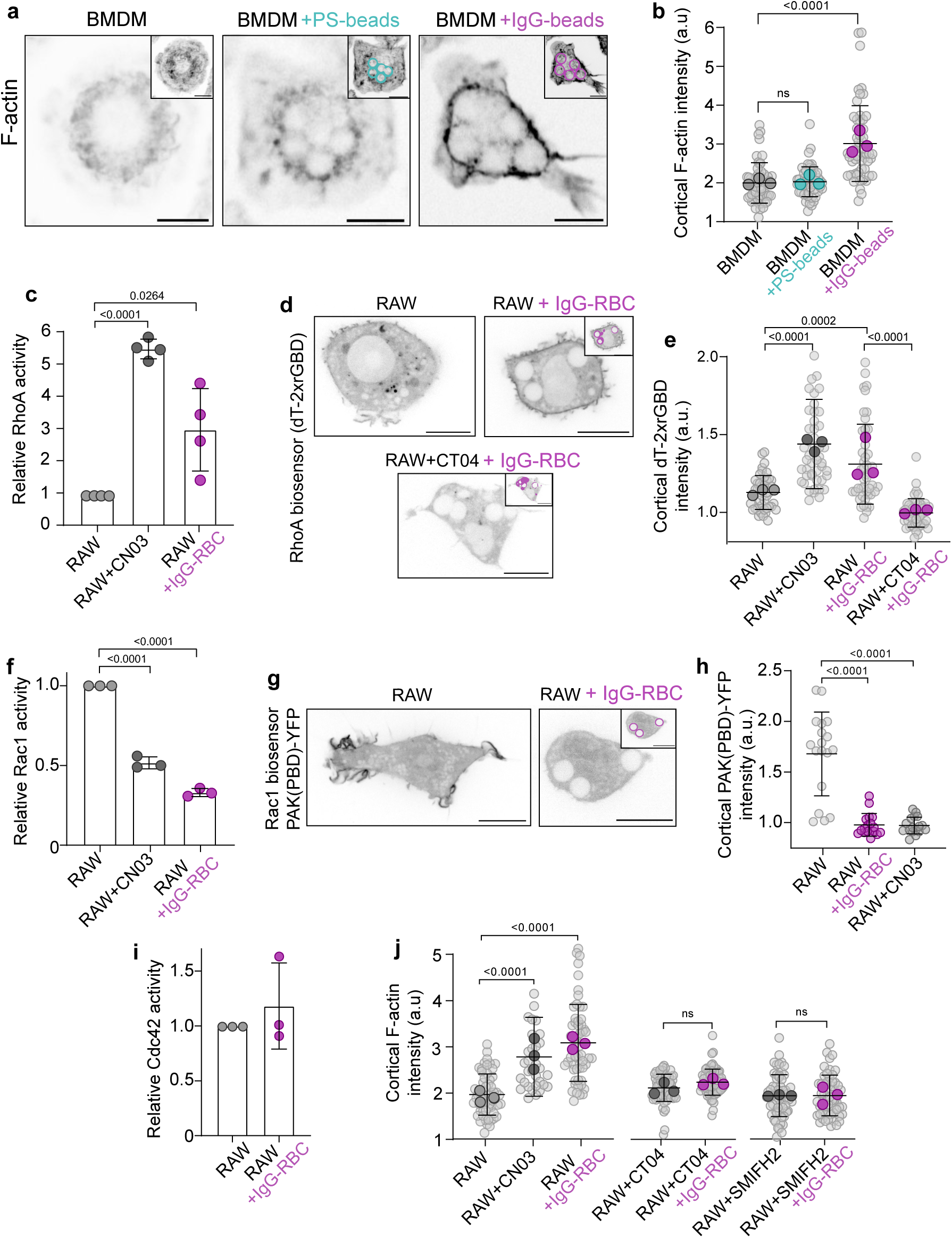
Cytoskeletal remodeling is associated with a sustained shift toward Rho/formin activity. **a-b)** In **a**, representative confocal images of the mid-section of BMDM stained with phalloidin (F-actin), 40 min after phagocytosis of PS-coated or IgG-opsonized beads. Insets are extended focus images showing PS-beads (cyan) or IgG-beads (magenta). In **b**, the normalized fluorescence intensity of the cortical F-actin was quantified in 3 independent experiments. **c, f, i)** Relative activity of indicated Rho GTPases measured by G-LISA in RAW 264.7 cells. In **c** and **f**, the cells were pretreated with the Rho activator, CN03, as specified in Methods. **d-e)** Confocal images of RAW 264.7 cells expressing dT-2xrGBD before and post-phagocytosis. In **e** the cortical dT-2xrGBD fluorescence, determined at a midsection plane, was normalized to the cytosolic fluorescence; data from 3 independent experiments. Where indicated, Rho was activated using CN03 or inactivated with CT04 as described in Methods. **g-h)** RAW 264.7 cells expressing PAK(PBD)-YFP before and after phagocytosis of indicated particles. In **h** the cortical PAK(PBD)-YFP signal was measured and normalized to the cytosolic fluorescence. **j)** Normalized cortical F-actin fluorescence intensity of RAW 264.7 cells containing ≥3 phagocytic targets in control cells or those treated with CN03, CT04, or the formin inhibitor SMIFH2 (5 μM); data from 3 independent experiments. *P* values are determined using a one-way ANOVA. All scale bars, 10 μm.

To determine if this feature was shared by other types of phagocytes, we also investigated the effect of IgG-opsonized particle uptake in human polymorphonuclear neutrophils (PMN). These cells display high motility, a process that is dependent on actin. We used video microscopy to track the motility of PMN before and after phagocytosis of IgG-opsonized red blood cells (RBC) (**Fig 1h-i, Video 2**). As shown in Fig 1h and quantified in Fig 1i, PMN the chemokinesis stimulated by N-formylmethionyl-leucyl-phenylalanine (fMLP) was virtually eliminated following engulfment of IgG-opsonized red blood cells (RBC). Therefore, a mechanism that arrests the surveillance programs upon ingestion of IgG-opsonized targets is conserved in various phagocytes.

### Phagocytosis of IgG-opsonized particles causes cortical thickening associated with increased activation of Rho

As described above, phagocytosis of particles via Fcγ receptors –but not PS receptors– induces major cytoskeletal rearrangements, a response that is proportional to the number of targets ingested (**S Fig 1**). The observed changes included a pronounced thickening of the submembranous F-actin (**Fig 2a-b**). Cortical F-actin dynamics are largely controlled by RhoA, which stimulates processive polymerization of actin filaments mediated by formins^14^. Accordingly, treatment of the macrophages with CN03^15^, a bacterial toxin-derived activator of Rho caused marked cortical thickening (not illustrated) that resembled the effects of Fcγ-mediated phagocytosis. Considering these similarities, we measured biochemically the activation of Rho following phagocytosis of IgG-coated particles. Because some of the agents used to manipulate and assess the activity of Rho interact with the A, B and/or C isoforms, we hereafter refer to their target as Rho(A-C). The effectiveness of the G-LISA assay used was validated using CN03, which induced a ≈5-fold stimulation of the GTPase. In good agreement with the microscopic observations of cortical F-actin, Rho(A-C) was activated nearly 3-fold following Fcγ receptor-mediated phagocytosis (**Fig 2c**).

While documenting its increased activity, the G-LISA assays do not reveal the site(s) where Rho is stimulated. Spatial information was obtained using a genetically-encoded biosensor of active Rho(A-C), namely dTomato-2xrGBD^16^, that contains the G protein-binding domain (GBD) of rhotekin, a Rho(A-C) effector. The biosensor was preferentially accumulated at the cortex of cells treated with CN03 and at the cortex of macrophages following phagocytosis of multiple IgG-coated particles. The redistribution of the dTomato-2xrGBD probe was prevented by CT04, a Rho(A-C) antagonist derived from *Clostridium botulinum* C3 transferase, validating the specificity of the probe (**Fig 2d-e**). These observations suggested that the activation of Rho(A-C) at or near the plasma membrane contributed to the cortical F-actin thickening described earlier.

The gain in Rho activity associated with phagocytosis was accompanied by a decrease in the activity of Rac^12^, whether assessed by G-LISA or using the PAK(PBD)-YFP biosensor^17^ (**Fig 2f-h**); the extension of ruffles where PAK(PBD)-YFP accumulates preferentially, was inhibited following phagocytosis of three or more IgG-coated particles. In contrast, we saw no appreciable difference in the overall activity of Cdc42 following phagocytosis (**Fig 2i**). To determine if the shift in the balance of GTPase activity towards Rho accounted for the cytoskeletal changes induced by phagocytosis, we quantified F-actin density at the cortex in RAW cells in the presence and absence of Rho(A-C) inhibitors. As in primary BMDM, RAW cells showed a clear increase in cortical F-actin post-phagocytosis of IgG-opsonized RBC to a level comparable to that observed following activation of Rho(A-C) with CN03 (**Fig 2j**). Conversely, inhibition of Rho(A-C) with CT04 prevented thickening of the cortex (**Fig 2j**).

Rho is anticipated to increase actin polymerization at the cortex by activating the formin family of nucleators that recruit actin monomers to growing linear filaments via their FH2 domains. RhoA, B, and/or C can activate mDia1, mDia2, Daam1, FMNL2, and FMNL3^18, 19^, though the relative contribution of individual formins to cortical F-actin is not clear. We therefore opted to broadly inhibit the formin family using the FH2 inhibitor (SMIFH2). As with inhibition of Rho itself, we found that inhibiting the formins also prevented thickening of the cortex in RAW cells that had ingested IgG-opsonized targets (**Fig 2j**). Taken together, these results implicate the gain in Rho(A-C) and formin activities in the cortical F-actin remodelling observed in macrophages containing phagosomes taken up via Fc receptors.

### The loss of podosomes associated with IgG-opsonized target engulfment is dependent on Rho/formin activation

In addition to cortex thickening, we previously noted that RAW cells that had ingested several IgG-opsonized targets lost the ventral F-actin puncta, reminiscent of podosomes, that are present in untreated cells. Immunostaining for the integrin-associated adaptor vinculin, confirmed that these structures were indeed podosomes (**Fig 3a**). Like the gain in cortical F-actin, the loss of podosomes was dependent on Rho(A-C) activation: inhibition by CT04 was sufficient to preserve the podosomes in RAW cells following ingestion of IgG-opsonized targets (**Fig 3b**). Reciprocally, the activation of Rho(A-C) alone, using CN03, eliminated the podosomes in otherwise untreated cells (**Fig 3b**). As was the case for the thickening of cortical F-actin, we found that formins were involved in the loss of podosomes, since inhibition with SMIFH2 prevented their disassembly following phagocytosis (**Fig 3b**). Taken together, these data implicated Rho/formins in the loss podosomes that accompanies the gain of cortical F-actin.

**Figure 3.**
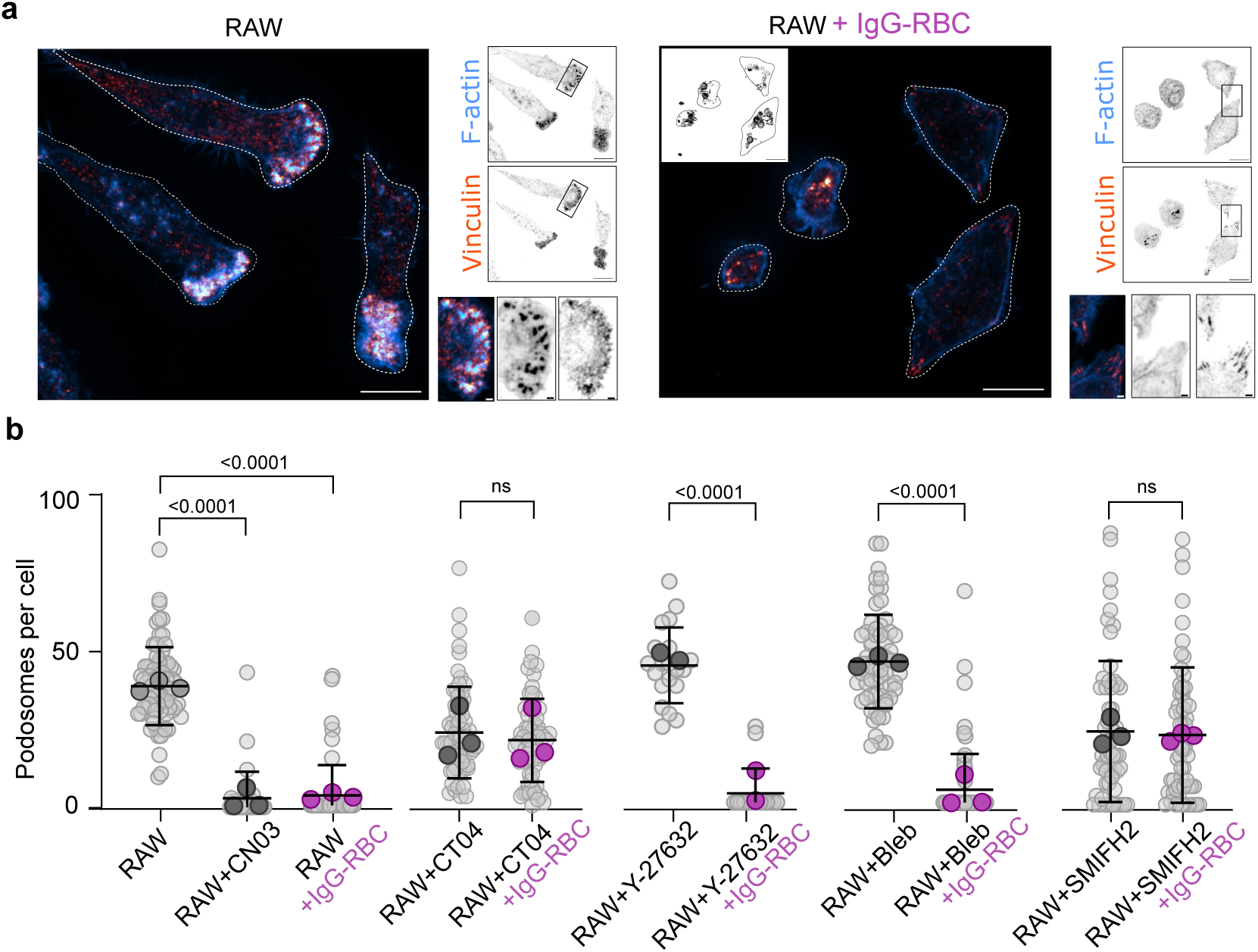
Loss of podosomes associated with Fc-mediated phagocytosis is the result of increased RhoA activation of formins. **a)** Ventral sections of RAW 264.7 cells before and 30-40 min after phagocytosis of IgG-RBC. Cells were stained for F-actin (blue) and vinculin (orange). Inset, projected image of particles. Scale bars, 10 μm. **b)** Quantification of the number of podosomes per RAW 264.7 cell before and after phagocytosis of >3 IgG-RBC particles. Cells were either untreated or pretreated with CN03 or CT04, then either pretreated with the ROCK inhibitor Y-27632 (5 μM) or with the myosin inhibitor blebbistatin (5 μM) for 10 min before exposure to phagocytic targets for an additional 30 min in the continued presence of the inhibitors. The formin inhibitor SMIFH2 (5 μM), which impairs phagocytosis, was only added after exposure and ingestion of phagocytic targets, then incubated an additional 30 min; data from 3 independent experiments. *P* values are determined using a one-way ANOVA.

Interestingly, when immunostaining for vinculin we noted that loss of podosomes was associated with a gain in focal adhesion-like structures (**Fig 3a**). Because the formation and signaling from focal adhesions are dependent on myosin II^20^, we suspected that the rounding and retraction of the cells following phagocytosis may involve this contractile protein. It is noteworthy that RhoA increases the activity of myosin II through its activation of Rho-associated kinase (ROCK), which phosphorylates myosin light chain. The involvement of these proteins can be tested experimentally, because ROCK and myosin II are potently inhibited by Y-27632 and blebbistatin, respectively. Neither of these inhibitors affected the binding or internalization of phagocytic targets (**S Fig 3**). Using these pharmacological agents we found that ROCK and myosin were in fact dispensable for the Rho-mediated loss of podosomes after Fc-mediated phagocytosis (**S Fig 3a-f**).

### ERM proteins are necessary to enact the changes in cell morphology associated with the engulfment of IgG-opsonized targets

It is remarkable that phagocytosis, which normally happens at or near the dorsal surface of the cells caused the loss of ventral podosomes. How this remote effect was exerted was not clear. It has been reported that changes in membrane tension can cause podosome disassembly^21, 22^, and the thickening or cortical actin that follows phagocytosis suggested that a similar process may be involved^23^. We therefore measured membrane tension by fluorescence lifetime imaging microscopy (FLIM) using the recently described Flipper probe^24^. As shown in (**Fig 4a**), membrane tension indeed increased as a result of phagocytosis, a change that persisted long after particle engulfment was completed (i.e. for at least 30 min). The tension change was mimicked by application of CN03, suggesting that cytoskeletal contraction due to stimulation of Rho(A-C) is responsible in both instances.

**Figure 4.**
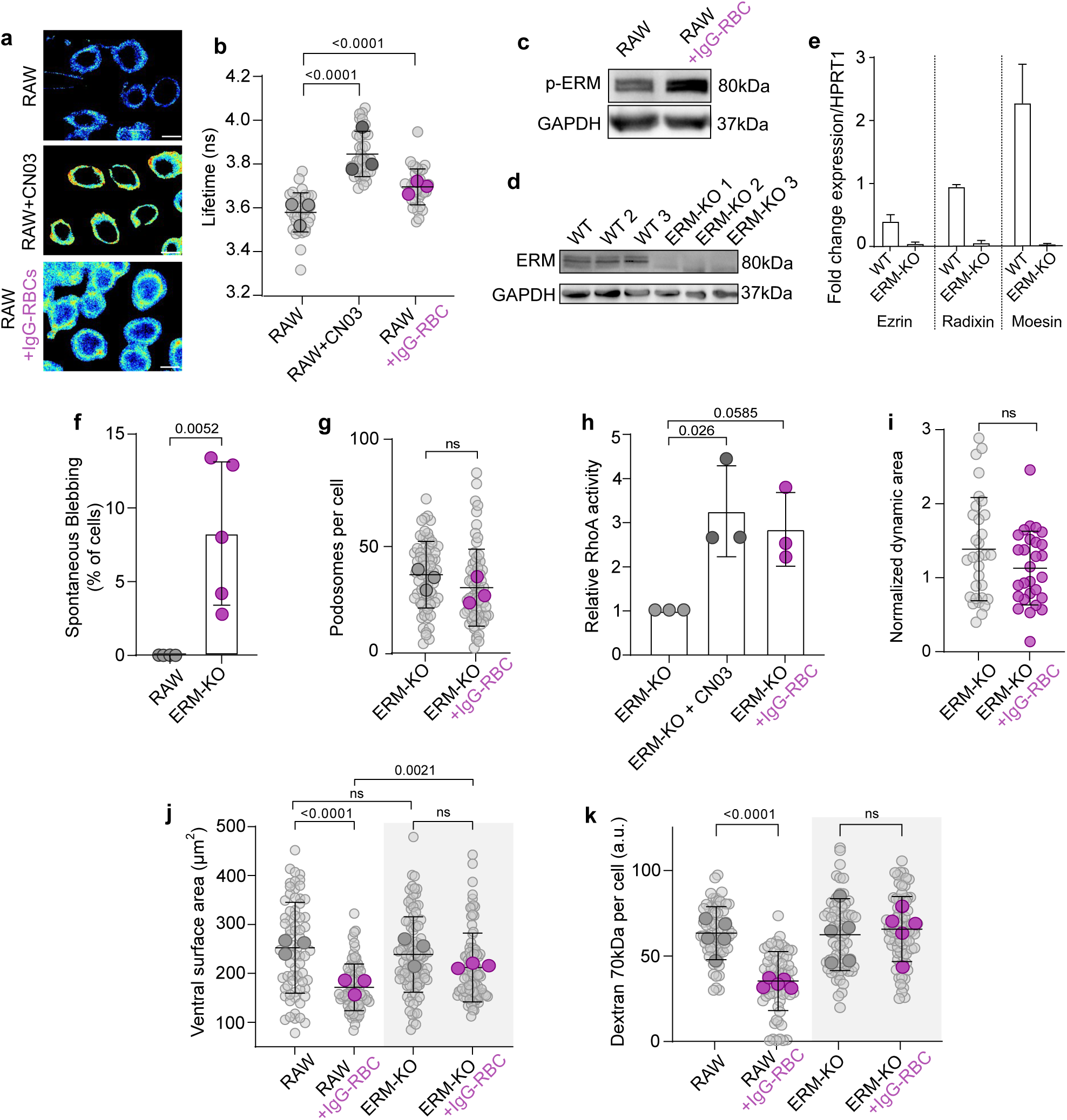
ERM proteins coordinate Rho/formin-mediated cortical cytoskeleton thickening, decreased membrane dynamics and podosome loss. **a-b)** Fluorescence lifetime imaging microscopy (FLIM) of FlipperTM in RAW 264.7 cells before and after phagocytosis of IgG-opsonized RBC. CN03-treated (Rho-activated) cells are also shown. Scale bar, 10 μm. The mean lifetime fluorescence determined in 3 separate experiments is graphed in **b**. **c)** Immunoblot of phospho-ERM (p-ERM) and GAPDH, used as loading control. Lysates were made 40 min after cells were challenged with particles. **d)** Immunoblot of the 3 independent clones where ezrin, radixin and moesin (ERM) were sequentially knocked out in RAW 264.7 cells. **e)** qPCR of ezrin, radixin and moesin in wildtype (WT) and ERM triple-KO cells. Data are means ± SD of 3 separate experiments. **f)** Spontaneous blebbing of triple-KO ERM (ERM-KO) in minimal medium. See also *S Video 3*. **g)** Quantification of podosomes per cell in ERM-KO cells before and 30-40 min after phagocytosis of IgG-opsonized RBC. Data from 3 independent experiments. Compare these values to those of Figure 3b. **h)** Rho activity measured by G-LISA in ERM-KO. Data are means ± SD from 3 separate experiments. **i)** Ruffling cell area of ERM-KO expressing Life-Act-GFP, determined as in Fig. 1e in, n = 3. See also *S Video 4*. **j)** Ventral surface area of ERM-KO and its parental RAW 264.7 cell line before and 40 min after phagocytosis of IgG-opsonized RBC. Surface was determined as in Fig. 1b. Only cells that contained 3 or more beads were quantified. Data from 3 independent experiments. **k)** TRITC-70 kDa dextran uptake after a pulse of 10 min by WT or ERM-KO cells before or after phagocytosis; data from 3 independent experiments. *P* values determined using a one-way ANOVA. In **f, g,** and **i,** a Student’s *t*-test was used.

We next considered how the compaction of the F-actin cytoskeleton exerted by Rho(A-C) activation is transmitted to the membrane. Such transmission would necessitate firm attachment of linear actin filaments to the membrane. The adaptor proteins ezrin, radixin and/or moesin (ERM) may fulfill such a role. ERM proteins are recruited by phosphatidylinositol-(4,5)- *bis*phosphate to the cytosolic leaflet of the plasma membrane, where they can then bind to multiple transmembrane proteins. Phosphorylation of ERM proteins near their C-terminus then facilitates their interaction with F-actin^25, 26^. We observed an increase in ERM phosphorylation following phagocytosis (**Fig. 4c**), supporting the idea that these proteins may contribute to the conveyance of tension to the membrane.

The possible role of ERM proteins in membrane tension and podosome disassembly was then assessed directly. Owing to their structural similarities, the three ERM proteins are believed to have overlapping functions^26^. Because RAW macrophages were found to express all three proteins, we were compelled to generate a triple ERM KO cell line using CRISPR-Cas9 editing. The elimination of the transcripts and resulting proteins was validated by PCR and immunoblotting, respectively (**Fig 4d-e**). Moreover, using single-particle tracking, we observed that CD44, a transmembrane protein that is anchored to the actin skeleton via ERM, displays a large increase in mobility in the KO cells^27^. As in a recent report^28^, we did not observe gross morphological differences between wildtype and ERM KO cells in normal culture medium. When incubated in minimal medium (HBSS), we did note a fraction of the ERM-KO cells underwent spontaneous blebbing, which was never observed in wildtype cells (**Fig. 4f** and **S Video 3**). Membrane blebbing became more pronounced upon activating Rho(A-C) with CN03 (**S Fig 4a-c**), supporting the idea that a decreased attachment of the membrane to the cortex increases blebbing when the cytoskeleton contracts^29^.

We proceeded to test whether ERM proteins are required for the cytoskeletal changes that follow Fc-mediated phagocytosis. Remarkably, we found that, unlike the wildtype cells, triple-KO ERM cells retained their podosomes after ingesting similar numbers of IgG-opsonized targets, despite undergoing a comparable increase in Rho activity (**Fig 4g-h**). We also found that the ERM KO cells continued to ruffle (**Fig 4i, Video 4**), performed macropinocytosis (**Fig 4k**), and remained more spread (**Fig 4j**) after phagocytosis than their wildtype counterparts. These data suggest that the concerted activation and function of Rho, formins, and ERM proteins are necessary to coordinate the phagocytosis-induced changes.

### The NADPH oxidase and associated ROS mediate the activation of Rho that renders the cells quiescent after phagocytosis

How does Rho(A-C) become activated downstream of Fc-receptor mediated phagocytosis? We initially considered the possibility that GEF-H1, a major Rho guanine nucleotide exchange factor (GEF) could be released into the cytosol following phagocytosis. GEF-H1 is made inactive by its association with microtubules and disassembly of the latter could account for its release and activation. However, we found that the microtubule cytoskeleton remained intact in the fed macrophages as determined by immunostaining and cytoskeletal fractionation (**Fig 5a-b**) suggesting this was not a likely mechanism of Rho activation.

**Figure 5.**
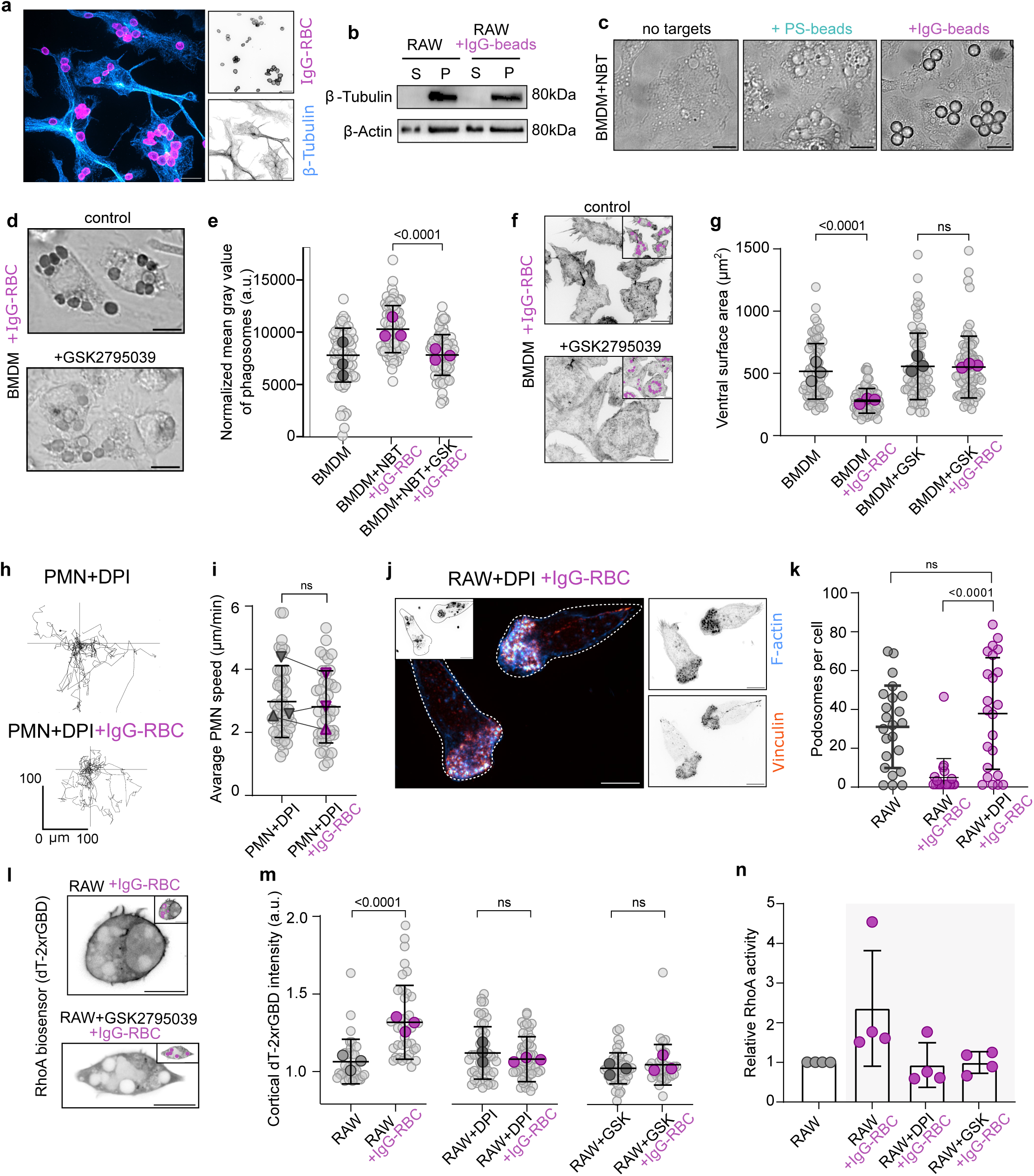
Reactive oxygen species generated during Fc-mediated phagocytosis trigger remodelling of the cytoskeleton. **a)** Immunostaining of β-tubulin (cyan) in BMDM 30-40 min after undergoing Fc-mediated phagocytosis. **b)** β-tubulin and β-actin associated with the pellet (P) after lysis in cytoskeletal-stabilizing buffer before and after phagocytosis, as indicated. **c)** Representative bright field image of BMDM 40 min after phagocytosis of either PS-coated or IgG-opsonized beads in the presence of nitroblue tetrazolium (NBT). **d)** Representative bright field image of BMDM 40 min after phagocytosis of IgG-opsonized RBC the presence of NBT with or without 5 μM of the NOX2 inhibitor, GSK2795039. **e)** Quantitation of the formazan deposited per phagosome. Data from 3 independent experiments. **f-g)** F-actin staining of BMDM 30-40 min post-phagocytosis of IgG-opsonized RBC in the presence or absence of 5 μM GSK2795039, added at the time of phagocytosis. Insets show phagocytic particles (magenta). In **g**, the ventral surface area of BMDM was quantified as in Fig. 1; n = 3. **h-i)** Chemokinesis of human neutrophils stimulated with fMLP before and after phagocytosis of IgG-RBC in the presence of diphenyleneiodonium (DPI; 10 µM). In **h**, trajectories of single cells from a representative video acquiring images every 10 s for 10 min. in **i** the average PMN speed was measured in 3 independent experiments. ▴ = female donor; ▾ = male donor. **j-k)** RAW264.7 cells treated with DPI and imaged 40 min after phagocytosis of IgG-opsonized RBC. Cells were fixed and stained for F-actin (blue), Vinculin (orange) and IgG (magenta). The number of podosomes per cell is shown in **k**. Dots represent individual cells from 3 independent experiments. **l-m)** RAW264.7 cells expressing dT-2xrGBD and treated with or without GSK2795039 were imaged after phagocytosis. The cortical dT-2xrGBD fluorescence intensity normalized to cytosol fluorescence is graphed in **m**. Data from 3 independent experiments. **n)** Rho activity in RAW264.7 cells measured by G-LISA under the indicated conditions. *P* values calculated by one-way ANOVA **(e, g, k, m)** or Student’s *t*-test **(i)**. All scale bars, 10 μm

A distinguishing feature between efferocytosis and Fc-mediated phagocytosis is the magnitude of the accompanying respiratory burst, which reflects the activity of the NADPH oxidase^30^. Upon incubating primary BMDM with phagocytic targets in the presence of nitroblue tetrazolium (NBT), formazan –a measure of reactive oxygen species (ROS)– was evident only in cells containing IgG-opsonized particles, but not PS-coated ones (**Fig 5c**). The inhibitor GSK2795039 prevented the response, confirming that the ROS were produced by NOX2, the predominant NADPH oxidase of phagocytes (**Fig 5d-e**). Of note, a number of studies have implicated ROS in regulating the activity of Rho proteins by oxidation of their GEFs and GAPs or of the GTPases themselves^31, 32^. The phagosome, like other membrane bilayers, is permeable to ROS so it seemed possible that oxidation of cytosolic determinants of Rho(A-C) activity could account for the morphological changes associated with Fc-mediated phagocytosis. Supporting this idea, inhibition of NOX2 with GSK2795039 prevented the rounding of the cells upon phagocytosis (**Fig 5g**). Moreover, diphenyleneiodonium (DPI), another potent inhibitor of the NADPH oxidase complex, prevented the arrest of chemokinesis in PMN following Fc-mediated phagocytosis (*compare* **Fig 5i** to **Fig 1i, Video 2** to **Video 5**). Lastly, podosomes persisted after phagocytosis when macrophages were treated with DPI (**Fig 5j-k**).

These data suggest a direct connection between the respiratory burst and the activation of Rho. We indeed noted that the targeting of the Rho biosensor to the cortex that follows phagocytosis in otherwise untreated cells was absent when the NADPH oxidase was inhibited (**Fig 5l-m**). We also found that the levels of active Rho were not elevated when the NADPH oxidase was inhibited by either DPI or by GSK2795039 (**Fig 5n**). Taken together, these data suggest that Fc-receptor triggered ROS production, mediated by NOX2, leads to Rho activation, causing changes in the shape and ability of phagocytes to migrate and survey their environment.

### Activation of the NADPH oxidase by pathogens and MAMPs drives podosome disassembly

In the preceding experiments we used non-biological targets to specifically contrast inflammatory vs. non-inflammatory responses. We next wondered how encounters with pathogens or soluble MAMPs would influence the surveillance activity of macrophages. Phagocytosis of pathogens generally triggers a respiratory burst as does signaling from TLR4^33^. We first challenged BMDM with heat-killed *Candida albicans*, an opportunistic pathogen that is bound and phagocytosed by receptors like the mannose receptor and Dectin-1 that recognize components of the fungal cell wall. After 40 min, we fixed the macrophages and subsequently stained for podosomal markers. As we had previously observed with IgG-opsonized targets, we found a remarkable change in the cell shape associated with the loss of podosomes and a concomitant gain in focal adhesion-like structures, illustrated in **Fig 6a** and quantified in **Fig 6b**. In addition, thickening of the cortical F-actin cytoskeleton was evident (**Fig 6c**). Therefore, the observations made using Fc-coated targets were recapitulated when stimulating fungal recognition receptors, which similarly activate the NADPH oxidase.

**Figure 6.**
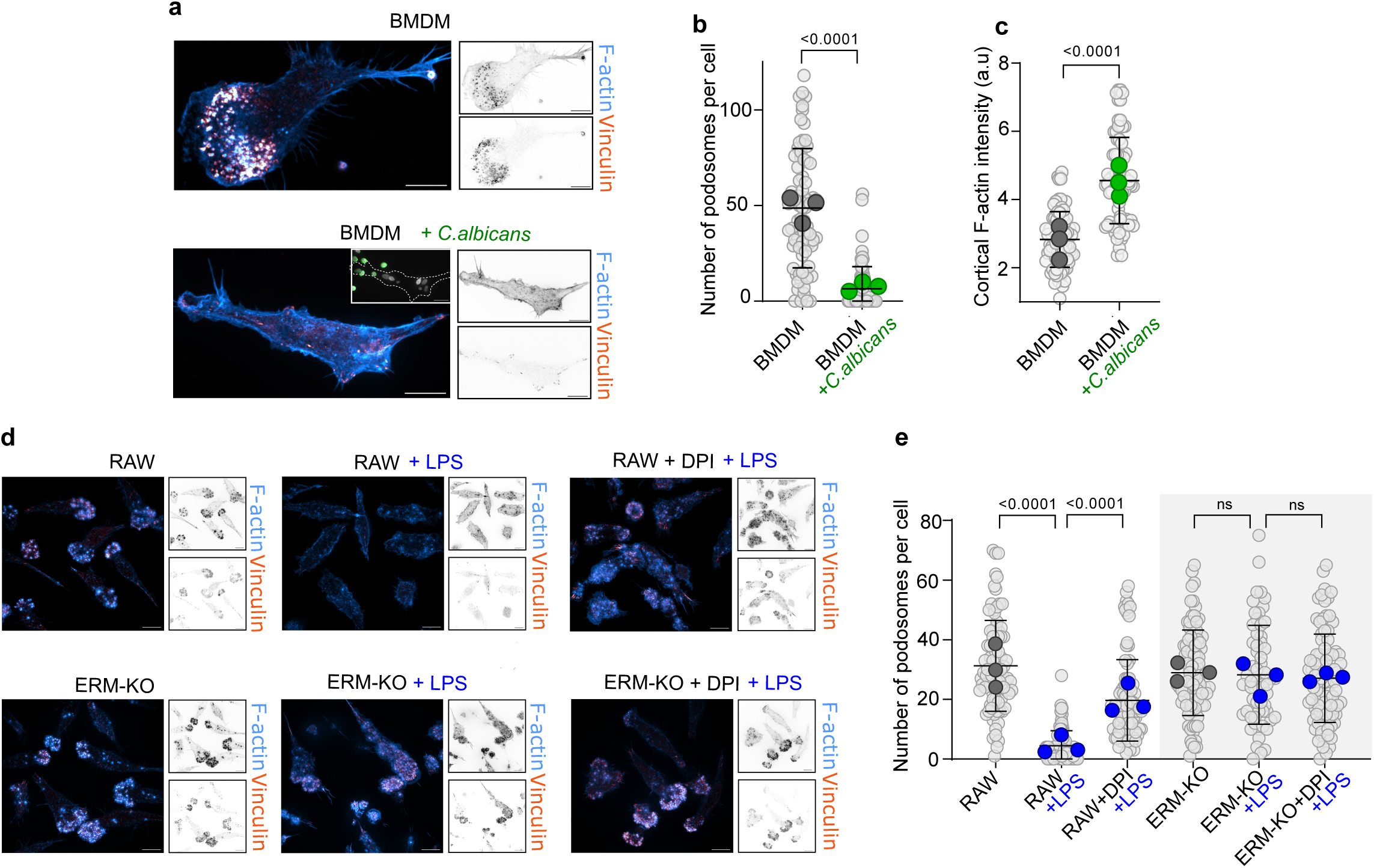
Pathogens and their associated molecules cause podosome loss by generating reactive oxygen species. **a)** BMDM differentiated in GM-CSF were incubated with or without heat-killed *C. albicans* expressing BFP (white). Cells were fixed and stained for concanavalin A to label extracellular *C. albicans* (green), then permeabilized and stained for F-actin (blue) and vinculin (orange). **b-c)** The number of podosomes per BMDM and the cortical F-actin intensity were determined as described in Fig. 1 in 3 independent experiments. **d-e)** Control wildtype or ERM-KO RAW 267.4 cells were treated with or without lipopolysaccharide (LPS) in the presence or absence of DPI, as indicated, then fixed and stained. In **e**, the number of podosomes per cell was determined in 3 independent experiments. *P* values are derived from one-way ANOVA **(e)** or Student’s *t*-test **(b-c)**. All scale bars, 10 μm.

The oxidase is also made active by LPS. Interestingly, exposure of cells to LPS had been previously shown to cause podosome disassembly^34^, which we were able to replicate (**Fig 6d-e**). While the mechanism of podosome loss in response to LPS is not entirely clear, it had previously been suggested that this could involve proteinases, notably the α-converting enzyme ADAM17. Alternatively, the cytoskeletal changes may be secondary to the activation of NOX2. We therefore investigated the role of *i)* the NAPDH oxidase and *ii)* ERM proteins in the disassembly of podosomes associated with TLR4 signaling. We found that inhibiting NOX2 caused the macrophages to retain their podosomes in LPS-treated cells, albeit with dysregulation of their positioning within the cells (**Fig 6d-e**). ERM proteins were also entirely necessary for the podosomes to disassemble in response to LPS (**Fig 6d-e**).

Taken together, these findings demonstrate that diverse signaling pathways downstream of pattern recognition receptors and Fc-receptors, which converge on the production of ROS via the NADPH oxidase, cause myeloid cells to change their phenotype from a motile to a quiescent state.

## Discussion

Myeloid cells continuously survey and sample their microenvironment; surveillance entails migration and extension of lamellipodia, while sampling involves specialized types of endocytosis that include macropinocytosis and phagocytosis. These processes all require dynamic remodelling of the cytoskeleton and have been observed to be antagonistic. Thus, patrolling of dendritic cells is arrested when they undergo a burst of macropinocytosis^35^. Changes in the symmetry of contractile F-actin networks are responsible for this transition^36, 37^. Migrating dendritic cells have myosin IIA-bundled filaments at their rear that propel the cell forward; upon macropinocytosis, these networks rapidly rearrange to become symmetric, temporarily preventing migration^37^. Global cytoskeletal rearrangements have also been reported to occur during phagocytosis. Elegant analyses in which phagocytes are held in place by a suction pipette while simultaneously exposing them to opsonized particles have revealed increases in cortical membrane tension at the back of the cell, as frontal pseudopods extend around the phagocytic target^38^. Similar results have been obtained by atomic force microscopy, measuring the rigidity of the dorsal membrane of macrophages undergoing frustrated phagocytosis^23^. While these mechanisms explain defective migration during the course of phagocytosis and macropinocytosis, the long-term consequences of particle engulfment had not been examined. As discussed earlier, the migratory behavior of the cells during the ensuing period of phagosome maturation and resolution has important consequences regarding the targeting of inflammation and the dissemination of infection.

Our studies revealed a prolonged quiescence, a state that extended for tens of minutes after the completion of Fc-mediated –but not PS-mediated– phagocytosis. A related effect had been observed 45 years ago for neutrophils: challenging the cells with IgG-coated beads impaired their chemotaxis through agarose^39^. More recent intravital studies have documented a complex behaviour of neutrophils described as swarming, which is characterized by chemotaxis followed by arrest at sites of tissue injury or infection^40^. The arrest has been attributed in part to desensitization of chemokine receptors, but cytoskeletal rearrangements resulting from phagocytosis may also contribute. For dendritic cells in the periphery, their migration towards the lymph nodes occurs much later (hours) after their phagocytosis of pathogens, in part owed to the need for their upregulation of chemokine receptors ^41^. Given our findings, the role of ROS in preventing dissemination, including via dendritic cells, should also be considered in orchestrating this sequence of events.

The Rho family of small GTPases are critical for coupling extracellular signals to rearrangements of the actin cytoskeleton^14^. In ostensibly unstimulated leukocytes the GTPases are kept in a critical balance that maintains spontaneous, dynamic surveillance. Our data suggest that ROS produced during phagocytosis can shift this balance to increase the activity of Rho(A-C) over that of Rac, arresting cell migration, ruffling, and macropinocytosis. Redox reactions have been described to modulate the function of a variety of cytoskeletal regulators, including the Rho GTPases themselves. RhoA is directly activated by ROS through oxidation of two cysteine residues (Cys16 and Cys20) located within the phosphoryl-binding loop of the GTPase. The redox-sensitive thiol of Cys20, in particular, is predicted to make direct contact with the bound nucleotide and increase nucleotide exchange; its oxidation has been shown to increase Rho activity as much as 600-fold^42^. The response of Rho GTPases to reactive species, however, is complex and varies among cell types. In muscle cells nitric oxide causes S-nitrosylation that inhibits RhoA, while peroxides and peroxynitrite stimulate RhoA activity in transformed and endothelial cells, respectively^43^.

It is noteworthy that various Rho-family GTPases, including Rac, possess a cysteine-containing motif (GXXXXGK[S/T]C) adjacent to their phosphoryl-binding loop^43^. Why then does Rho(A-C) activation prevail over Rac in phagocytes that have activated the respiratory burst? The location of the various Rho GTPases vis-à-vis the phagosomal membrane, where most of the ROS are generated, may be of relevance. Rac is a direct activator of the NADPH oxidase so would be, in principle, expected to be susceptible to oxidation^30^. Interestingly, Rac-GTP directly associates with other enzymes that regulate redox homeostasis, including the superoxide dismutase, SOD1^44^. Also, Rac may be negatively regulated by oxidation of its Cys178 that would prevent its modification by palmitic acid, required for membrane recruitment^45^. Redox reactions extend also to regulators of Rho GTPases. Exposure of endothelial cells to peroxynitrite or peroxide, for example, increases protein kinase C (PKC)-mediated phosphorylation and activation of p115-RhoGEF, which leads to increased levels of RhoA-GTP^46^. A respiratory burst produced by the NADPH oxidase may therefore result in the (reversible) activation of Rho over Rac via multiple direct or indirect convergent mechanisms.

In phagocytes, the net effect of stimulated RhoA activity is the inhibition of migration, ruffling, and podosome formation, all resulting from increased polymerization of linear actin filaments via formins. These filaments furnish and thicken the cortex of the cell, providing sites for actin-binding proteins to bridge transmembrane proteins to the cortical skeleton. Supporting this model, we found that inhibiting formins or reducing the connections between membrane proteins and submembranous actin filaments by deleting ERM proteins promoted sustained formation of podosomes and ruffling of the cells. Conversely, stimulating ERM activity would be expected to counteract cellular surveillance. ERM proteins need to be phosphorylated to become fully active, and we noted an increase in the phosphorylation of ERMs that was sustained long after Fc-mediated phagocytosis. Interestingly, PKC, which is oxidation-sensitive, as well as Rho kinase (ROCK), are responsible for this phosphorylation^26^. This would suggest exquisite synchronization between the polymerization of linear actin networks and their attachment to the plasma membrane when cells mount a respiratory burst. Together, these responses drive the cells into a quiescent state with increased cortical rigidity and a predicted decrease in the diffusion of membrane proteins.

Rho activation occurs also during mitosis, in that instance as a result of RhoGEF1 release from microtubules that depolymerize during cell division. Interestingly, ERM proteins are also phosphorylated and made more active in mitosis, an event that is important for the reported increase in cortical membrane tension in dividing cells. In *Drosophila*, which express only moesin, its deletion alone causes defects in cytokinesis, with failure to increase cortical tension^47^. After mitosis, the basal architecture of the actin cytoskeleton is re-established, enabling cells to regain their primary shape and attachment to the substratum. The parallel changes observed during mitosis and following phagocytosis indicate that similar means, i.e. activation of Rho(A-C), are used in various settings to enact marked changes in cellular structure and function.

Based on the results of this study, we posit that phagocytic cells gauge pathogenic threats by oxidizing self-molecules via the products of the NADPH oxidase. The assembly and activation of the oxidase are regulated by several processes, notably the phosphorylation of its components, their binding to phospholipids, and the recruitment and stimulation of Rac2^30, 48^. While the activation of the complex is envisaged to occur early in phagocytosis, the duration of the stimulated state has not been clearly established and reinvigoration of the complex at late stages may well occur if infectious particles persist or even grow within the host cell. Regardless of the precise timing when Rho is activated to exert its inhibitory effects on cell motility, this mechanism constrains inflammation and limits the dissemination of pathogens, which could otherwise employ migrating immune cells as Trojan horses to enter distant tissues.

## Materials and Methods

### Cell lines and cell culture

RAW 264.7 cells were from ATCC. These cells were cultured at 37°C in an atmosphere of 95% air and 5% CO_2_ in DMEM (Wisent Inc.) supplemented with 10% heat-inactivated fetal bovine serum (FBS; Gibco). Unless otherwise indicated, the cells were plated sparsely on glass coverslips within 12-well tissue culture plates (Corning Inc.) and allowed to grow overnight.

BMDM were grown out from the long bones of 6-12-week-old C57BL/6 wild-type mice isolated as previously described^49^. Briefly, bones from euthanized mice were dissected, immersed in 70% ethanol for 5 min, dried, and the ends removed. The bone marrow was extruded by centrifugation (15,000 g, 10 s) into cold phosphate-buffered saline (PBS) and after removal of the bones the cells were pelleted (500 g, 10 min). Pellets were suspended in DMEM with 10% FBS, 10 ng/mL M-CSF, 1× antibiotic-antimycotic solution, and plated at a density of 4.0 × 10^5^ cells per 10 cm Petri dish for 5–8 days before use. For experiments measuring podosome formation, BMDM were suspended in DMEM with 10% FBS, 10 ng/mL gm-CSF, 1×antibiotic-antimycotic solution, and plated at a density of 8.0 × 10^5^ cells per 6-well dish before use.

For human neutrophils, all reagents underwent filtration through Detoxi-Gel Endotoxin Removing Gel Columns (Thermo Fisher Scientific) before the isolation. Human neutrophils were isolated utilizing a density-gradient separation method as described previously^50^. Briefly, 30 mL of blood was aseptically collected into BD Vacutainer EDTA tubes (BD Biosciences, Mississauga, ON, Canada) and then gently layered on top of PolymorphPrep (Progen, Wayne, PA, USA) at a 1:1 ratio. The layered suspension was subsequently centrifuged at 500 g with an acceleration of 1 and deceleration of 0 for 30 min to effectively separate neutrophils from mononuclear cells. The isolated neutrophils were carefully washed with Hank’s balanced salt solution supplemented with calcium and magnesium (Wisent). The resulting neutrophils were finally resuspended in HBSS and utilized within 2 h for all experimental procedures.

The *Candida albicans* clinical isolate SC5314-BFP was grown in YPD (BD Difco™ YPD) at 37°C overnight, then heat-killed at 65°C, washed and used for phagocytosis experiments.

### Reagents

Anti-rabbit secondary antibodies conjugated with Alexa Fluor 488 and Alexa Fluor 647 were from Jackson ImmunoResearch Labs. Triton X-100 was from Fisher Scientific, Acti-stain was from Cytoskeleton, Inc. Protease inhibitor cocktail was from Pierce. Lifeact-EGFP was sourced from Dyche Mullins and obtained as Addgene plasmid #58470, while dTomato-2xrGBD was acquired as Addgene plasmid #129625.

### Rho Activation and Inhibition Experiments

Rho Activator II (Cytoskeleton, Cat# CN03) and Rho Inhibitor I (Cytoskeleton, Cat# CT04) were utilized on the same day after dilution. Alternatively, these reagents were stored at - 80°C for up to 2 weeks. Cells were grown overnight in DMEM supplemented with 10% FBS, the medium was then replaced with HEPES-buffered Tyrode’s solution and incubated for 10 min before the addition of the CN03 or CT04. HEPES-buffered Tyrode’s Solution contained: HEPES: 25 mM, D-Glucose: 10 mM, KCl 5 mM, NaCl 140 mM, MgCl_2_ 1 mM, CaCl_2_ 2 mM, in Autoclaved MilliQ Water (H_2_O), adjusted to pH 7.4. The cells were incubated for 1 h at 37°C in an incubator without CO_2_. After incubation, the culture medium was exchanged with complete medium containing 10% serum also supplemented with the peptides. The cells were further incubated for 1 h at 37°C in a 5% CO_2_ environment before performing experiments.

### Transfection

RAW 264.7 cells were transfected using FuGene HD (Cat# E2311, Promega) following the manufacturer’s instructions. 6 h after transfection, the medium was replaced with fresh culture medium.

### Generation of ERM-knockout cells

To generate moesin-knockout cells, RAW264.7 cells were transfected with a CRISPR gRNA plasmid DNA (U6-gRNA:CMV-Cas-9-2A-tGFP) from Sigma-Aldrich, with a specific target sequence for moesin (5’-CCGGCTTCGGATTAACAAG-3’), using FuGENE HD. 24 h post-transfection, cells expressing GFP were sorted by FACS and individually plated in 96-well plates to establish single-cell colonies. Individual colonies were expanded, and the absence of moesin expression was verified by immunoblotting using the Q480 monoclonal antibody (Cat# 3150S, Cell Signaling). To generate double EM knockouts, the procedure was repeated transfecting moesin KO cells with the ezrin target sequence 5’-TGTGGCACGCGGAACACCG’. The absence of ezrin expression was confirmed through immunoblotting with an anti-ezrin antibody (Cat# 3145S, Cell Signaling). Triple ERM knockouts were generated using the radixin target region sequence was 5’-AAAACAGTTGGCTTACGTG’, and radixin expression was confirmed by immunoblotting using the monoclonal C4G7 antibody (Cat# 2636S, Cell Signaling). Negative control cell lines, which retained normal expression of ERMs, were selected from the same population of cells used to isolate the ezrin and radixin knockouts. ERM knockouts and negative controls underwent final validation steps through immunoblotting using an antibody against ezrin/radixin/moesin (Cat# 3142S, Cell Signaling) and by qPCR.

### Western blotting

Cell lysates were prepared using cold RIPA buffer from Sigma supplemented with EDTA-free protease and phosphatase inhibitors from Thermo. Cell lysates were cleared by centrifugation at 13,000 g for 15 min at 4°C. Protein concentration was determined using the BCA assay. Laemmli buffer (BioRad) containing β-mercaptoethanol (Sigma) was added to the lysates, and the samples were boiled at 95°C for 5 min. These prepared lysates were then loaded onto 10% SDS-PAGE gels. Following separation in the SDS-PAGE gels, the samples were transferred to PVDF membranes using the Bio-Rad wet transfer system. The transferred membrane was subsequently blocked with a solution of 5% bovine serum albumin (BioShop) in Tris-buffered saline (TBS) for 1 h at room temperature. The membrane was then subjected to overnight incubation at 4°C with primary antibodies, which were diluted in blocking buffer. Primary antibodies used included: a 1:1000 β-tubulin antibody from Abcam (Cat# ab15568), a 1:1000 β-actin antibody from Abcam (Cat# ab8227), a 1:1000 ezrin/moesin/radixin antibody from Cell Signaling (Cat # 3142), and 0.4 μg/mL GAPDH from Millipore (Cat# MAB374). After the primary antibody incubation, membranes were washed 4 x 5 min with TBS containing 0.1% Tween-20 (TBST). Subsequently, the membranes were incubated for 2 h at room temperature with appropriate secondary antibodies (Jackson ImmunoResearch) diluted in blocking buffer. The secondary antibodies used were 1:3000 horse radish peroxidase (HRP)-conjugated donkey anti-mouse antibody from and 1:3000 HRP-conjugated donkey anti-rabbit antibody. Following the secondary antibody incubation, the membranes were washed 4 x 10 min with TBST. Chemiluminescence signals were developed using ECL^TM^ Prime Western Blotting Detection Reagents from Cytiva and detected using the ChemiDoc XRS System from BioRad. For the microtubule precipitation experiments, we used a microtubule stabilization buffer containing: PIPES 80 mM, MgCl_2_ 2 mM, EGTA 1 mM, GTP 1 mM, glycerol 50%, Triton X 0.5%, supplemented with protease and phosphatase inhibitors.

### G-LISA

RAW 264.7 cells were grown overnight in 6-well plates. Cells were challenged with IgG-opsonized RBC for 40 min. Rho-GTP was measured using the RhoA G-LISA Activation Assay Kit (Colorimetric Format; Cat# BK124) from Cytoskeleton, according to the manufacturer’s protocol.

### Microscopy

Confocal images were captured using a spinning disk system (WaveFX; Quorum Technologies Inc.), consisting of a microscope (Axiovert 200 M; Zeiss), a scanning unit (CSU10; Yokogawa Electric Corporation), an electron-multiplied charge-coupled device camera (C9100-13; Hamamatsu Photonics), a five-line (405 nm, 443 nm, 491 nm, 561 nm, and 655 nm) laser module (Spectral Applied Research), and a filter wheel (MAC5000; Ludl). The system is controlled by Volocity software version 6.3. Confocal images were obtained using a 63×/1.4 numerical aperture oil objective from Zeiss, coupled with an additional 1.53 magnifying lens, and the appropriate emission filters. For live-cell imaging, glass coverslips were mounted within a Chamlide magnetic chamber (Live Cell Instrument Inc.) and overlaid with pre-warmed HBSS. The temperature was maintained at 37°C using an environmental chamber (Live Cell Instruments Inc.).

### Fluorescence staining

Paraformaldehyde (4% wt/vol)-fixed cells were permeabilized in 0.1% (vol/vol) Triton X-100 in PBS for 10 min, blocked in 2% (wt/vol) BSA in PBS for 40 min and overlaid consecutively for 2 h with primary and secondary antibodies in 2% BSA, separated by PBS and BSA washes. Phagocytic efficiency was assessed by differential inside–outside staining.

### FLIM (Fluorescence Lifetime Imaging Microscopy)

Cells were incubated with 1 µM Flipper-TR for 5-15 min before imaging. FLIM measurements in the frequency domain were conducted using an Olympus IX81 inverted microscope equipped with a Lambert-FLIM attachment. This setup included a ×60/1.49 NA oil immersion objective and a Li2CAM iCCD camera. The modulation frequency was set at 400 MHz, and the instrument was calibrated based on the known fluorescence lifetime of Alexa Fluor 546, assuming a monoexponential lifetime of 4.1 ns. The acquired lifetime data was subsequently analyzed using FLIM software from Lambert Instruments.

### Image analysis and statistics

Image handling, quantification, and analysis of fluorescence images were performed using Volocity 6.3 (PerkinElmer Inc), Imaris 9.5.1 (Oxford Instruments), or ImageJ 1.54f (NIH).

### Statistical analysis

Prism (GraphPad) was used to perform all statistical analyses. For comparison between two groups, unpaired two-tailed Student’s t-test was used. For comparisons among multiple groups one-way ANOVA with Tukey’s multiple comparisons test was used. Unless otherwise stated, error bars are representative of mean ± standard deviation of the mean (SD). P values smaller than 0.05 were considered statistically significant.

## Notes

### Competing Interest Statement

The authors have declared no competing interest.

### Summary of Updates

Discussion has been expanded and some references have been added

