## Supplementary Figures for "Reactive oxygen species suppress phagocyte surveillance by oxidizing cytoskeletal regulators"

### Supplementary Figure 1

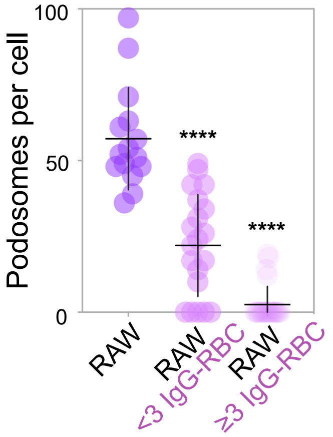


SF1. Quantification of the number of podosomes per RAW 264.7 cell before and after phagocytosis of <3 IgG-RBC and ≥3 IgG-RBC.

### Supplementary Figure 2

#### SF2. Flowchart of procedure used for quantification of dynamic area in RAW 264.7 cells

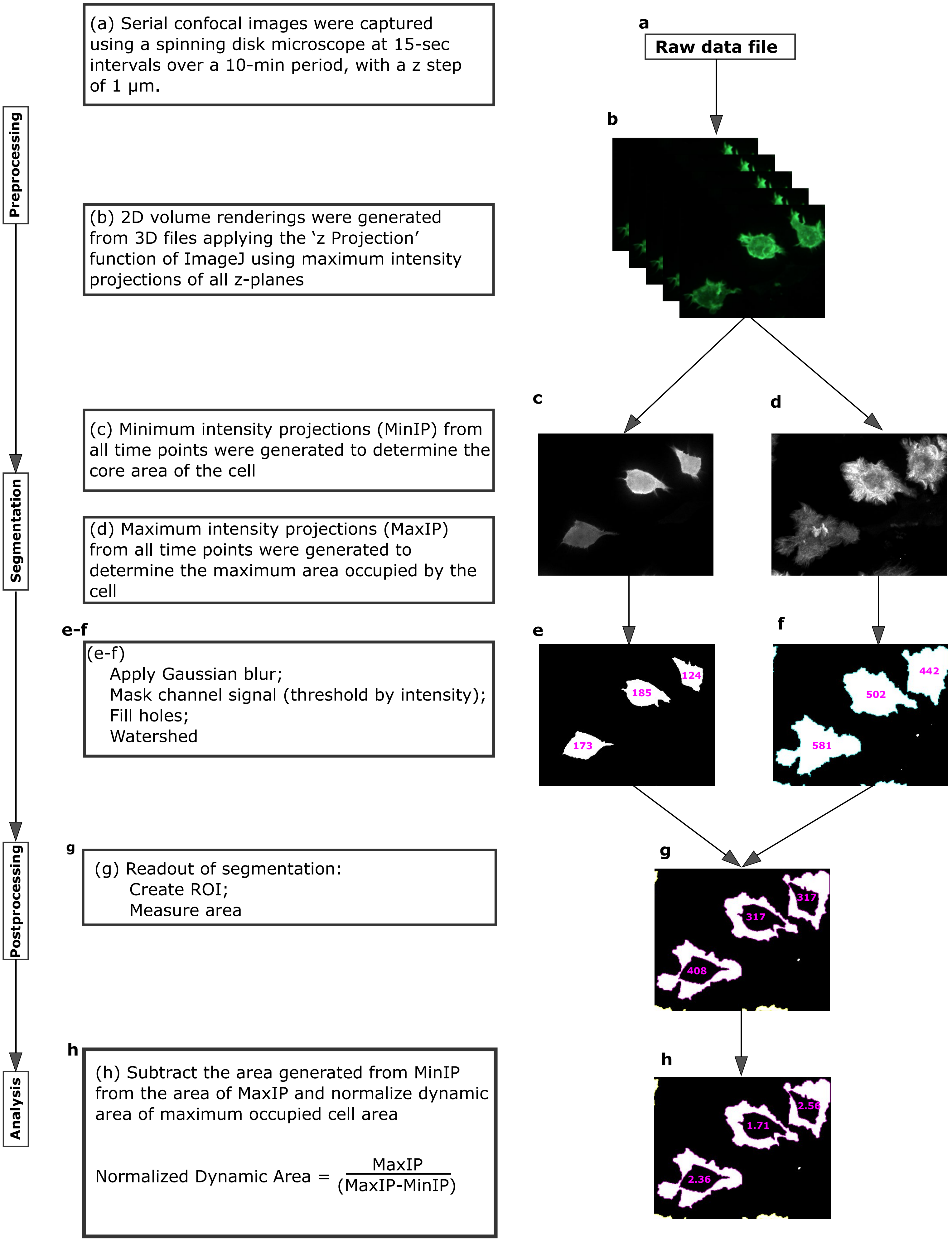

#### Supplementary Figure 3

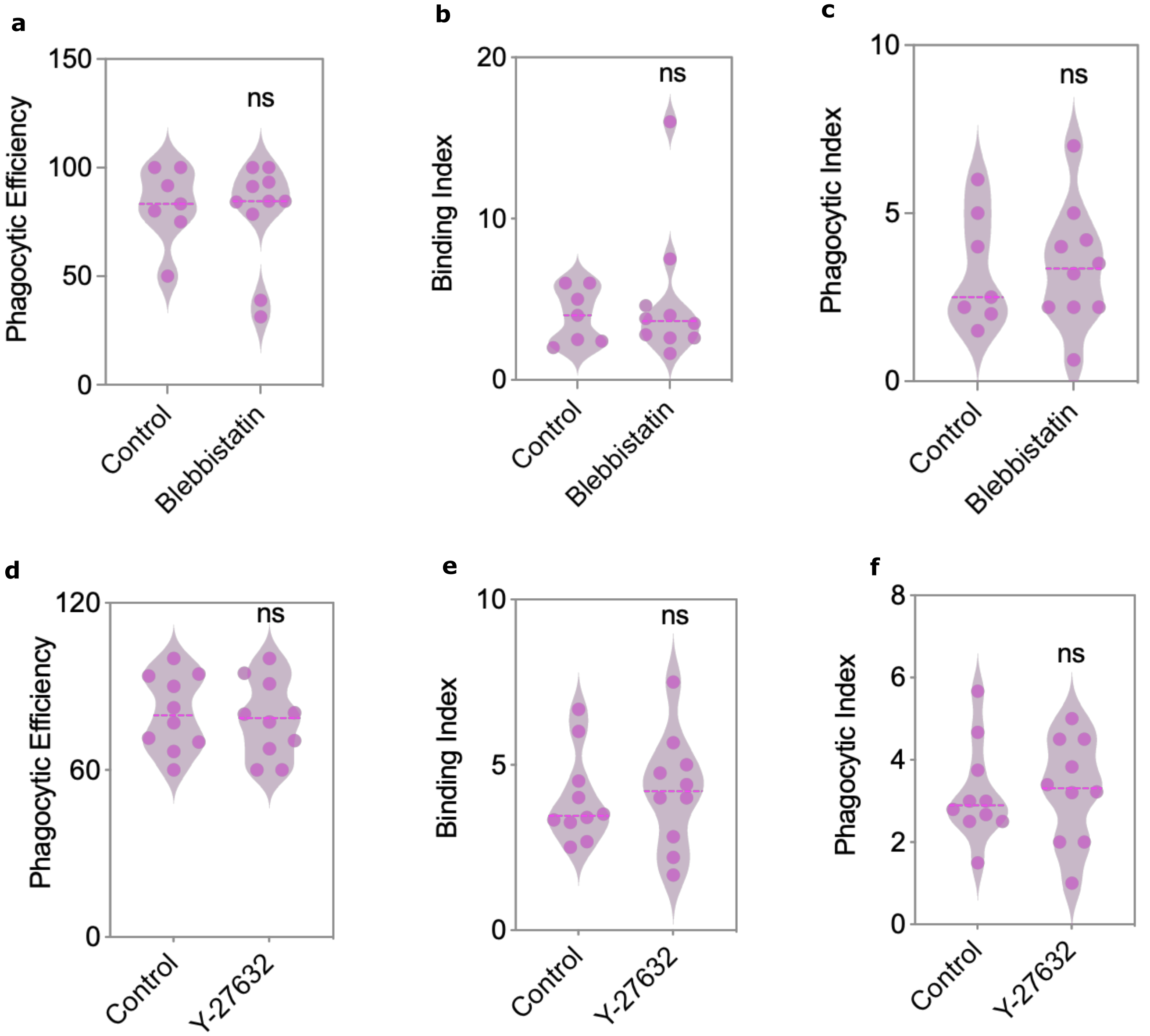


SF3. **(a, b, c)** Violin plots showing phagocytic efficiency, binding index and phagocytic index of RAW 264.7 cells that were otherwise untreated (control) or were pretreated with 5 μM blebbistatin for 10 min before and also during phagocytosis. Each dot represents a separate field of view, cells from 3 independent experiments. **(d, e, f)** Violin plots showing phagocytic efficiency, binding index and phagocytic index of RAW 264.7 cells that were otherwise untreated (control) or were pretreated with ROCK inhibitor 5 μM Y-27632for 10 min before and also during phagocytosis.. Each dot represents a separate field of view, cells from 3 independent experiments

#### Supplementary Figure 4

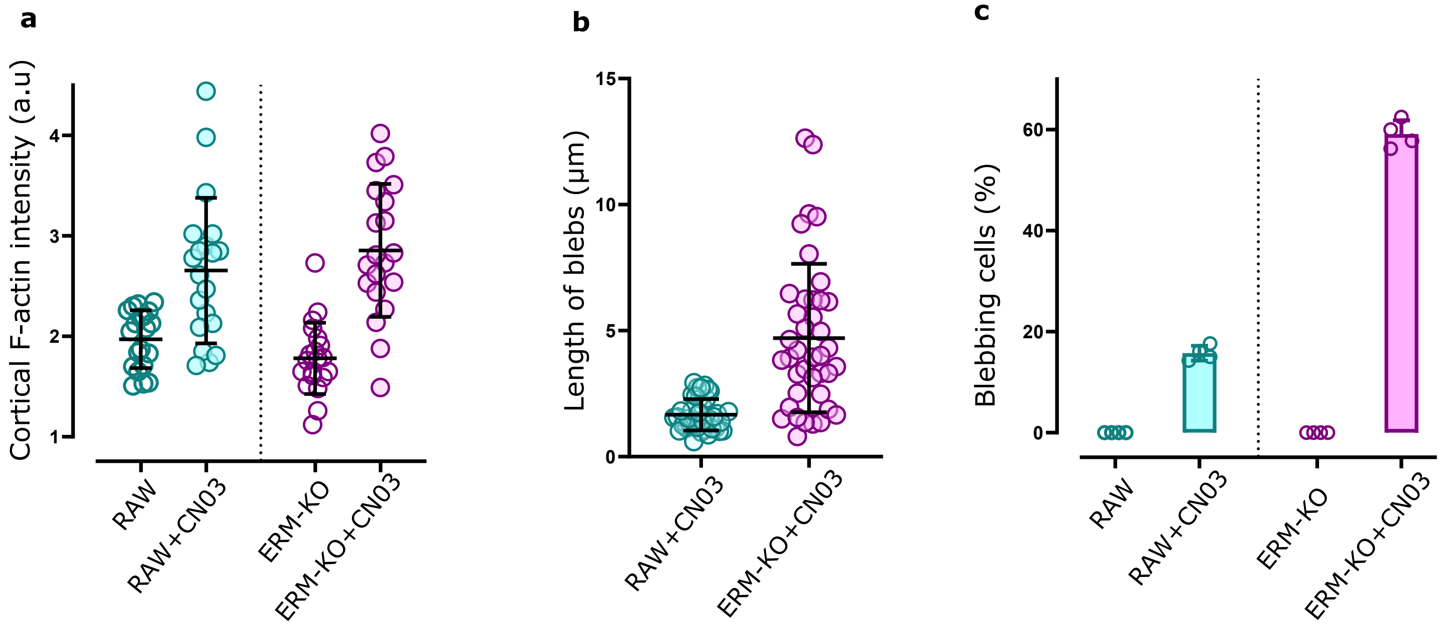


SF4. **(a)** Quantification of normalized fluorescence intensity of cortical F-actin measured at the mid-section of RAW264.7 and ERM-KO cells before and after CN03 treatment. Dots represent individual cells. **(b)** Length (in µm) of blebs appearing in RAW264.7 and ERM-KO cells after CN03 treatment. **(c)** Quantification of the percentage of blebbing cells; RAW264.7 and ERM-KO cells were bathed in RPMI medium supplemented with 10% FBS before and after CN03 treatment.  Each point represents an independent experiment
